# Inhibition of Multiple Staphylococcal Growth States by a Small Molecule that Disrupts Membrane Fluidity and Voltage

**DOI:** 10.1101/2024.01.17.576101

**Authors:** Jamie L. Dombach, Grace L. Christensen, Samual C. Allgood, Joaquin LJ Quintana, Corrella S. Detweiler

**Author notes:** The views expressed in this presentation/manuscript are those of the authors and do not necessarily reflect the official policy of the Department of Defense, Department of Army, US Army Medical Department, or the US Government.

## Abstract

New molecular approaches to disrupting bacterial infections are needed. The bacterial cell membrane is an essential structure with diverse potential lipid and protein targets for antimicrobials. While rapid lysis of the bacterial cell membrane kills bacteria, lytic compounds are generally toxic to whole animals. In contrast, compounds that subtly damage the bacterial cell membrane could disable a microbe, facilitating pathogen clearance by the immune system with limited compound toxicity. A previously described small molecule, D66, terminates *Salmonella enterica* serotype Typhimurium (*S.* Typhimurium) infection of macrophages and reduces tissue colonization in mice. The compound dissipates bacterial inner membrane voltage without rapid cell lysis under broth conditions that permeabilize the outer membrane or disable efflux pumps. In standard media, the cell envelope protects Gram-negative bacteria from D66. We evaluated the activity of D66 in Gram-positive bacteria because their distinct envelope structure, specifically the absence of an outer membrane, could facilitate mechanism of action studies. We observed that D66 inhibited Gram-positive bacterial cell growth, rapidly increased *Staphylococcus aureus* membrane fluidity, and disrupted membrane voltage while barrier function remained intact. The compound also prevented planktonic staphylococcus from forming biofilms and disturbed three-dimensional structure in one-day-old biofilms. D66 furthermore reduced the survival of staphylococcal persister cells and of intracellular *S. aureus*. These data indicate that staphylococcal cells in multiple growth states germane to infection are susceptible to changes in lipid packing and membrane conductivity. Thus, agents that subtly damage bacterial cell membranes could have utility in preventing or treating disease.

**Importance:** An underutilized potential antibacterial target is the cell membrane, which supports or associates with approximately half of bacterial proteins and has a phospholipid makeup distinct from mammalian cell membranes. Previously, an experimental small molecule, D66, was shown to subtly damage Gram-negative bacterial cell membranes and to disrupt infection of mammalian cells. Here we show that D66 increases the fluidity of Gram-positive bacterial cell membranes, dissipates membrane voltage, and inhibits the human pathogen *Staphylococcus aureus* in several infection-relevant growth states. Thus, compounds that cause membrane damage without lysing cells could be useful for mitigating infections caused by *S. aureus*.

## Introduction

New clinical antibiotics are often chemical variants on natural products that microbes have deployed as antimicrobial weapons for millions of years. Bacteria quickly evolve resistance to chemical analogs of existing antibiotics by modifying endogenous pathways or by acquiring new resistance genes horizontally (1, 2). Moreover, during infection, bacteria are frequently differentially susceptible to antimicrobials because they adopt different growth states, including planktonic, biofilm, and persister states (3, 4). Therefore, compounds that interfere with infection by disrupting new targets across multiple growth states are needed.

Sources of new potential antimicrobials include screening platforms based on cell culture infection models, which recapitulate specific aspects of host soluble innate immunity better than traditional screens in standard microbiological media (5, 6). Multiple research groups have therefore developed screening platforms for compounds that prevent the survival of bacterial pathogens in the phagolysosomes of macrophages, which are professional phagocytes of the monocyte lineage that reside within tissues. In-macrophage screens for anti-infectives have utilized bacterial pathogens such as *Salmonella enterica* and *Mycobacterium tuberculosis* (7–10). These works have identified compounds that lack antibacterial activity in standard microbiological media but inhibit bacterial growth under broth conditions that mimic the macrophage phagosome environment. For instance, such compounds prevent *S.* Typhimurium replication in broth in growth media or altered genetic backgrounds that increase outer membrane permeability and/or inactivate efflux pumps (7, 9, 11–14). Thus, the relative impermeability of the Gram-negative cell envelope in standard media complicates study of the mechanism of action for these compounds.

The present work focuses on the mechanism of action and the effect on disease-relevant growth-states of the small molecule D66 (378.3 g/mol) in Gram-positive bacteria because they lack an outer membrane barrier. D66 is a screening hit from a commercially available drug-like discovery library (7, 15). The compound is minimally toxic to mammalian cells and mice, prevents the replication of *S.* Typhimurium in macrophages, and curbs infection in mice. Under broth conditions that permeabilize the outer membrane or reduce efflux pump activity, D66 inhibits the growth of Gram-negative bacteria and disrupts voltage across the inner membrane (11). In Gram-positive bacteria, we found that exposure to the compound inhibited growth, rapidly increased membrane fluidity, and disrupted membrane voltage without permeabilizing cells. D66 exposure inhibited biofilm formation and dispersed one-day old biofilms. The compound also reduced the survival of staphylococcal persister cells in broth and in mammalian cells. These data suggest that compounds that disturb but do not readily lyse membranes have the potential to interfere with multiple bacterial growth states relevant to infection.

## Results

### Growth of Staphylococcus species and Bacillus subtilis is inhibited by D66

To establish whether Gram-positive bacteria are sensitive to D66, we monitored bacterial growth across a dose range of D66 in four staphylococcal strains and in a *B. subtilis* laboratory strain. The origins and defining characteristics of these strains are described in the methods. Bacterial growth (absorbance) over 18 hours was monitored after dilution of overnight cultures to an OD_600_ of 0.01 in LB containing D66. All strains tested were unable to grow in the presence of high concentrations of the compound, with values of half minimum inhibitory concentrations (MIC_50_) ranging from 43 - 93 µM (**Fig 1 A, B**). In contrast, *S.* Typhimurium with an intact outer membrane grew normally even at 150 µM of D66, a concentration approaching the limit of solubility (11). Thus, bacteria lacking an outer membrane barrier have increased susceptibility to D66 upon out-growth from overnight cultures.

**Figure 1.**
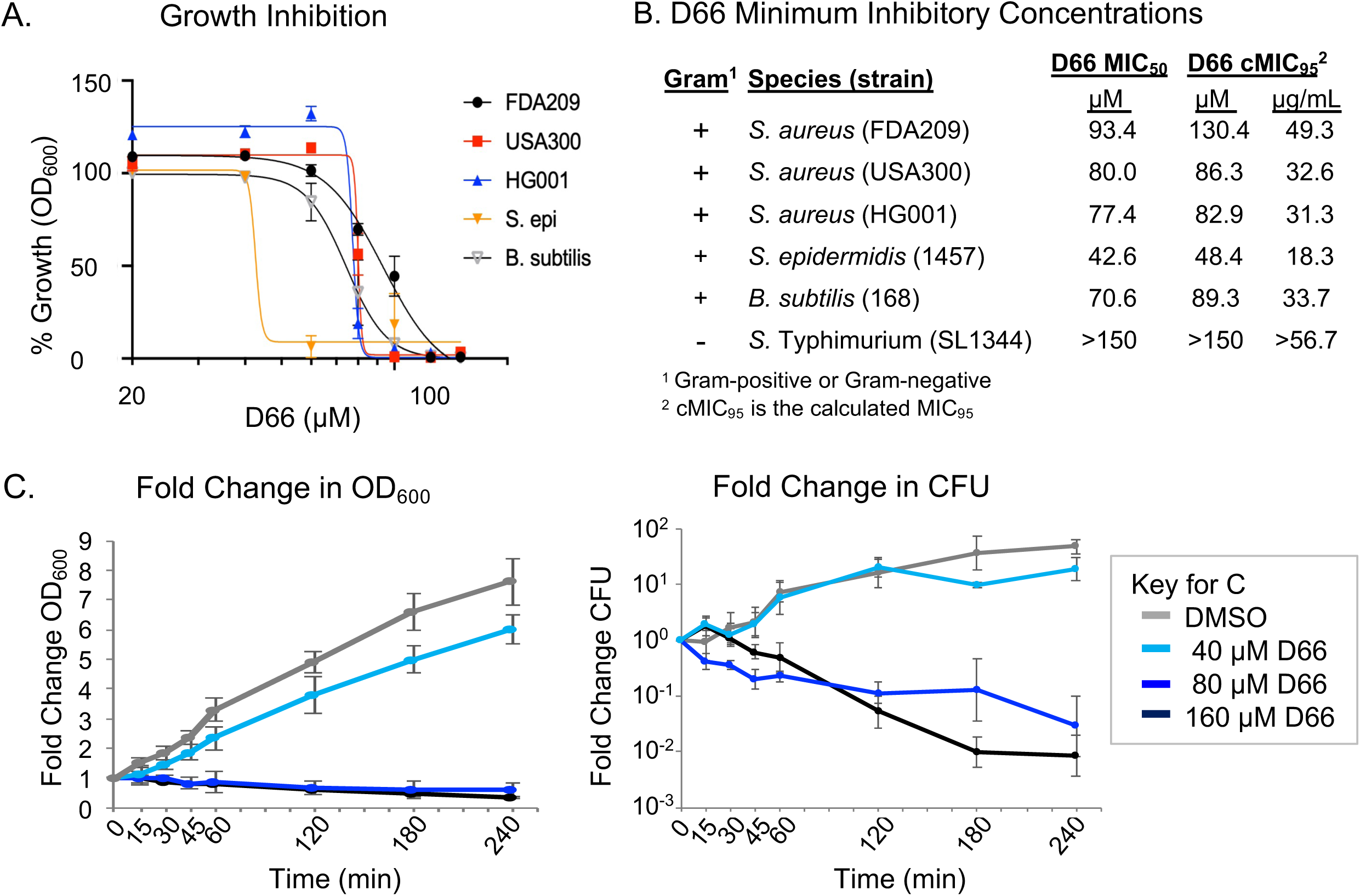
D66 inhibits the growth of Gram-positive bacteria in broth and is bactericidal to mid-log phase *S. aureus*. **A)** Dose response curves monitoring staphylococcal and *B.* subtilis percent growth inhibition in the presence of D66. Data are normalized to growth in 2% DMSO after 18 hours. Mean and SEM of at least three biological replicates performed with technical triplicates. **B)** Table of D66 concentrations that inhibit growth*. S.* Typhimurium is included for comparison. **C)** Fold change in OD_600_ and CFU/mL. Mid-log phase cultures of *S. aureus* FDA209 were treated at time 0 with DMSO or D66. Concentrations of D66 above and below the MIC_50_ and cMIC_95_, ranging from 40 – 160 μM were tested. Cultures were monitored for absorbance and plated for enumeration of CFU over 4 hours. Data are presented as fold change. Mean and SEM of three biological replicates performed with technical triplicates.

We evaluated whether D66 could affect bacterial survival in mid-log phase cells. *S. aureus* FDA209 was exposed to three concentrations of compound, and growth and survival (colony forming units, CFU) were monitored over four hours (**Fig 1 C**). At 40 μM of D66, strain FDA209 grew slightly less than DMSO control treated cells. At 80 μM of D66, no growth was observed, and absorbance decreased over time, suggesting bacterial lysis. These observations were validated by the 10-100-fold decline in CFU from the initial cell count. We conclude that D66 is bactericidal to *S. aureus* in mid-log phase, and that *S. aureus* could be used in experiments aimed at understanding D66 activity across bacterial growth states germane to infection.

### S. aureus cell membranes rapidly increase fluidity in the presence of D66

Prior work in Gram-negative bacteria suggested the bacterial cell membrane is a target of D66 (11). Therefore, we established whether D66 could affect critical aspects of staphylococcal cell membrane function, including fluidity. Changes in membrane fluidity are detectable with the fluorescent probe laurdan, which inserts into the phospholipid bilayer. The emission spectrum of laurdan changes according to the polarity of the environment, as determined by calculating the generalized polarization (GP) (16–18). Treatment with the membrane fluidizer benzyl alcohol (BnOH) rapidly decreased GP, as expected (12, 19) (**Fig 2 A**). Exposure to 40 μM D66 similarly decreased GP, indicating membranes become more fluid in the presence of this compound. At 160 μM of D66, membrane fluidity quickly reversed, and rigidity increased, potentially reflecting D66 self-interactions. D66 thus rapidly increases *S. aureus* cell membrane fluidity.

**Figure 2.**
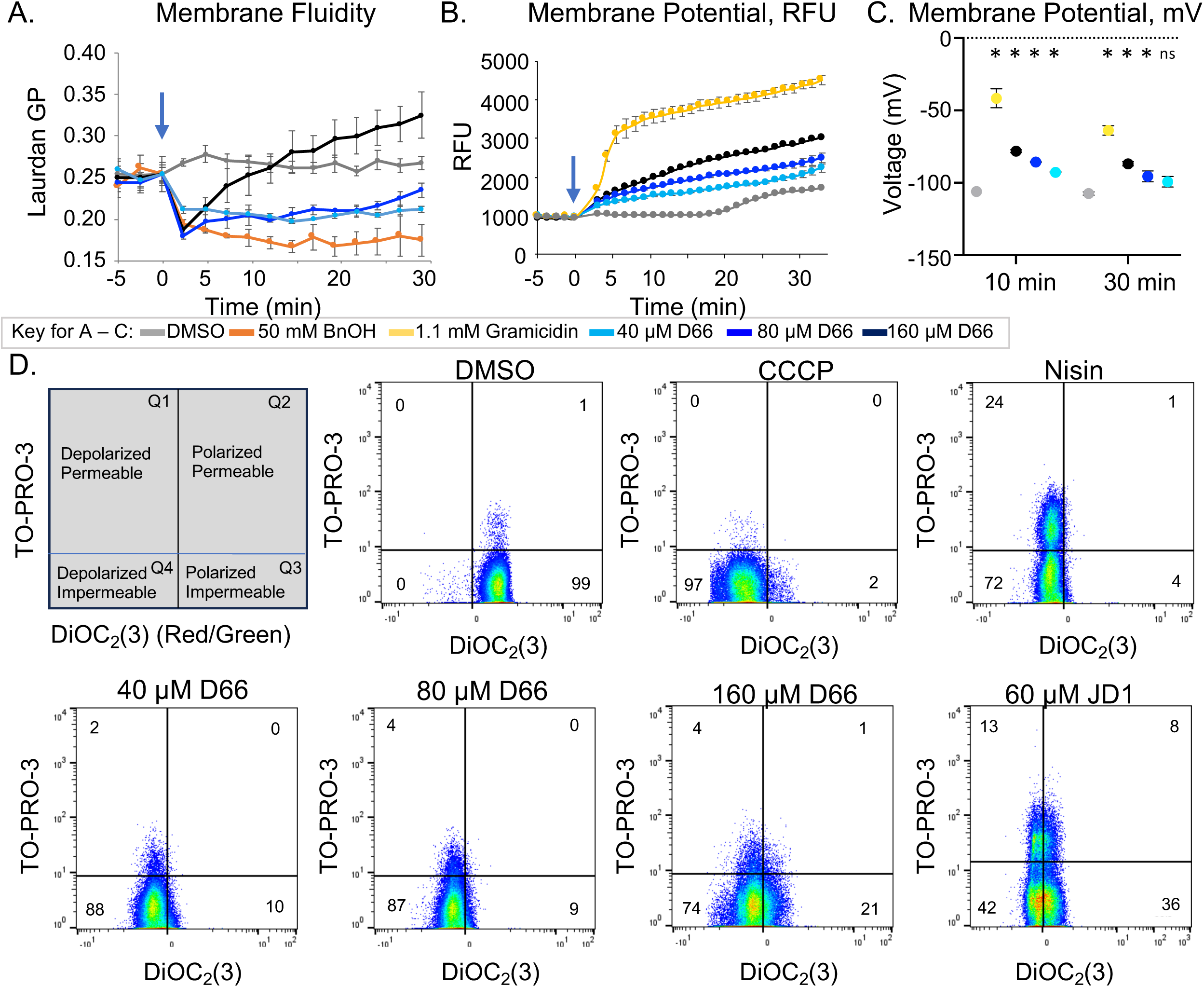
*S. aureus* cell membranes become more fluid and depolarize upon exposure to D66 but are not significantly permeabilized. Mid-log phase *S. aureus* FDA209 cells were used for all experiments. **A)** Membrane fluidity was evaluated with a Laurdan generalized polarization (GP) assay. Cells were treated just after time 0 (arrow) with DMSO, the membrane fluidizer benzyl alcohol (BnOH) [50 mM], or D66. Mean and SEM of three biological replicates performed with technical triplicates normalized to DMSO at time 0. **B)** Membrane voltage was monitored with the fluorescent dye DiSC3(5) and reported as relative fluorescent units (RFU). Cells were treated just after time 0 (arrow) with DMSO, the ionophore gramicidin (2 μg/ml; 1.1 mM) or D66. Mean and SEM of three biological replicates performed with technical triplicates normalized to DMSO at time 0. **C)** Membrane potential in mVolts was calculated at 10- and 30-minutes after treatment based on calibration curves (Figure S1B). Asterisks indicate *P* ≤ 0.01 compared to the corresponding DMSO sample, determined by one-way ANOVA with a Dunnett’s post-test; ns, not significant. **D)** Membrane polarization and permeability were determined by flow cytometry using the fluorescent dyes TO-PRO-3 and DiOC_2_(3). Cells were treated with DMSO, CCCP [30 μM], nisin (75 μg/mL), D66 or JD1 for 5 minutes. Data are representative of three biological replicates, with percentages are shown in each quadrant.

### D66 disrupts membrane voltage without creating large holes

Increases in membrane fluidity can impact ion and proton distribution across the cell membrane (20–22). To monitor membrane voltage upon exposure to D66 we used the fluorescent probe 3,3’-dipropylthiadicarbo-cyanine iodide (DiSC3(5)), which is quenched upon intercalation into charged lipid bilayers. Voltage disruption releases DiSC3(5) from the membrane and increases fluorescence (23, 24). Treatment of bacteria with gramicidin pore forming peptides [1 mM] depolarized membranes and increased DiSC3(5) fluorescence, as expected (25), changing voltage from approximately -106 mV to -42 mV within 10 minutes (**Fig 2 B, C, Fig S1A**). Treatment with D66 modestly increased DiSC3(5) signal, for instance, changing voltage to -93 mV and -78 within 10 minutes at, respectively, 40 and 160 μM. In summary, D66 disrupts membrane voltage in a dose responsive manner.

Changes in membrane fluidity and voltage disruption can reflect decreased membrane barrier function (16, 18, 26). However, we did not observe significant changes in the proton gradient, reduction potential, nor the ability to exclude propidium iodide upon exposure to D66 until at least 30 minutes after treatment at the highest dose (**Fig S1 B - E**). Since these assays were conducted across populations of bacteria, we turned to flow cytometry to determine whether, in individual cells, membrane permeability and potential change simultaneously upon treatment with D66. We used the fluorescent dyes 3,3′-diethyloxacarbocyanine iodide (DiOC_2_(3)) and TO-PRO-3 to determine whether D66 exposure depolarizes cells and/or compromises membrane integrity, respectively. DiOC_2_(3) accumulates in membranes and fluoresces red if the membrane is polarized. DiOC_2_(3) fluorescence shifts to green if voltage decays; the red/green fluorescence ratio provides a measurement of depolarization. If the cell membrane is permeable, TO-PRO-3 diffuses into the nucleoid and binds DNA, increasing its fluorescence. Treatment of *S. aureus* with CCCP decreased the red/green DiOC_2_(3) fluorescence ratio without affecting TO-PRO-3 signal, as expected (**Fig. 2 D, Fig S1 F - H**). Nisin, a small antimicrobial peptide that forms pores in the membrane, increased TO-PRO-3 fluorescence and decreased the DiOC_2_(3) fluorescence ratio, as anticipated. JD1, a small molecule that lyses bacterial membranes (27) demonstrated increased TO-PRO-3 signal. Treatment of the bacteria with D66 decreased the DiOC_2_(3) fluorescence ratio with only a marginal increase in TO-PRO-3 fluorescence, up to 11 +/- 3% within 45 minutes at the highest concentrations of D66. These observations indicate that in most cells D66 rapidly disrupts membrane voltage without creating large holes. Overall, the data suggest that D66 drives increases in membrane fluidity that perturb voltage, but does not lyse cells.

### D66 inhibits biofilm formation in the *S. aureus* HG001 strain

When bacteria transition from a planktonic growth state to a biofilm, their cell membranes undergo a decrease in fluidity (28–31). Therefore, we investigated whether exposure of planktonic cells to D66 affects biofilm formation. Staphylococcus strains were exposed to a range of sub-growth-inhibitory concentrations of D66 over 24 hours, and the accumulation of extracellular mass was monitored with crystal violet staining (**Fig 3 A-D**). Rifampin (rifampicin) and vancomycin were selected as positive control antibiotics because they are effective against planktonic *S. aureus* and diffuse through biofilms (32, 33). Crystal violet staining revealed that in strain FDA209, only rifampin inhibited biofilm formation. In HG001 and *S. epidermis,* the 1/2x MIC concentrations of rifampin and D66 (respectively, 0.025 μg/mL;30 nM and 40 μM) reduced biofilm formation. The USA300 strain was prone to increased biofilm formation, especially at lower concentrations of rifampin, vancomycin, and D66. To establish whether D66 differentially affects the accumulation of live or dead cells in the maturing biofilm, we imaged live biofilms with a spinning disc confocal microscope. Relative levels of live (Syto9, green) and dead (PI, red) cells were quantified. Vancomycin exposure increased the volume of live and dead cells in the HG001 strain, mirroring observations in other vancomycin-sensitive *S. aureus* strains (34). In contrast, both rifampin and D66 reduced the volume of live and dead cells in HG001 (**Fig 3 E**). In the *S. epidermidis* strain, rifampin reduced cell volume, and D66 increased the volume of live and dead cells (**Fig 3 F**). Thus, D66 exposure has mixed effects but inhibits biofilm formation in the HG001 strain nearly as well as rifampin by reducing the volume of live and dead cells.

**Figure 3.**
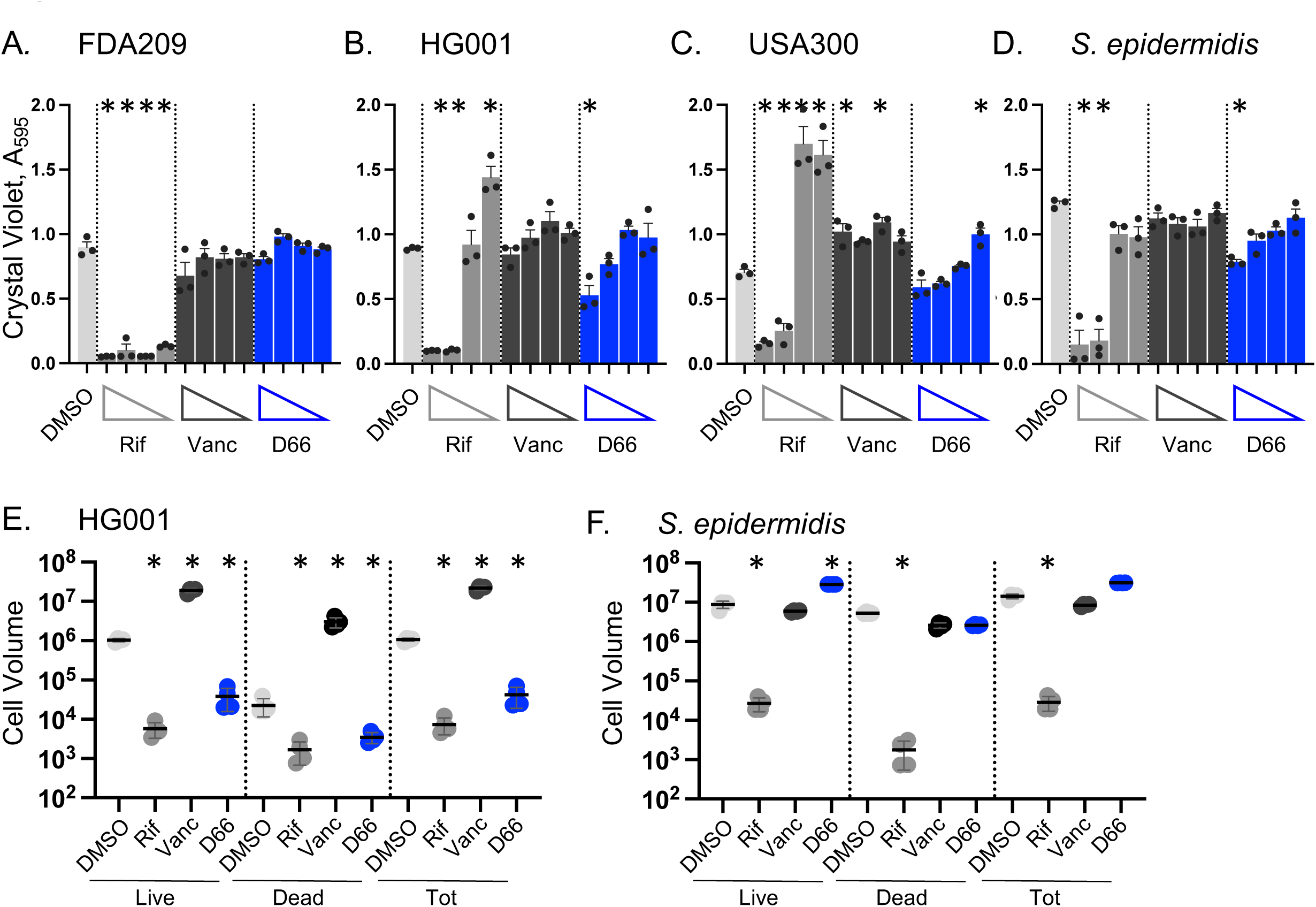
D66 has modest effects on biofilm formation in staphylococcus. Biofilm formation was monitored after 18 hours in TSB with subinhibitory concentrations of DMSO, rifampin (Rif; 1x MIC_95_ = 0.05 µg/mL), vancomycin (Vanc; 1x MIC_95_ = 1 µg/mL), or D66 (1x MIC_95_ = 80 µM). **A - D**). Crystal violet staining for accumulated biofilm mass measured at A_595_. Concentrations of Rif, Vanc and D66 were ½ x, ¼ x, 1/8 x and 1/16 x x MIC_95_ from left to right. Mean and SEM of three biological replicates performed with technical triplicates and normalized to untreated cells. Asterisks indicate *P* ≤ 0.05 compared to DMSO, determined by a one-way ANOVA with a Dunnett’s post-test. **E, F**). Biofilms incubated with ½ x MIC of the indicated compound were stained with Syto 9 (live cells) and PI (damaged/dead cells) and imaged with confocal fluorescent microscopy. Volume of live, dead, and total cell volume was calculated (positive volume = positive voxel; 1 voxel = 0.0367 µm3). Left, Cell Volume; Right, ratio of live/dead cells. Mean and SD of one of two biological replicates derived from a minimum of four fields of view. Asterisks indicate *P* ≤ 0.01 compared to DMSO, determined by a one-way ANOVA with a Tukey post-test on log_10_ transformed data.

### In one-day-old biofilms, D66 reduces the number of live cells and disrupts biofilm architecture

We next examined whether D66 disrupts the maintenance of 1- and 5-day old biofilms. Biofilms were treated with DMSO, rifampin, vancomycin, or D66 for 18 hours. The mass of one-day-old biofilms, as measured by crystal violet, was reduced in the FDA209 *S. aureus* and the *S. epidermidis* strains in response to rifampin. Vancomycin treatment only affected the *S. epidermidis* strain (**Fig 4 A-D**). In contrast, D66 reduced one-day-old biofilm mass in the FDA209 and USA300 strains. Rifampin and D66 at 4x MIC disrupted biofilm structure and reduced the volume of live cells in *S. aureus* HG001 and in *S. epidermidis* (**Fig 4 E, F**). Volume reconstructions of the imaged biofilms confirmed that after one day, *S. aureus* HG001 formed thinner (∼38 µm) biofilms than *S. epidermidis* (∼80 µm) (4, 35). Exposure to D66 at 4x MIC reduced biofilm height and complexity similarly to rifampin (**Fig 4 G, H**). In 5-day old biofilms, vancomycin had little effect on any strain, but rifampin and D66 reduced mass in HG001 (**Fig S1**). In summary, D66 treatment reduces the number of live cells in HG001 and in *S. epidermidis* one-day-old biofilms. D66 also diminishes biofilm structure, suggesting that compounds that alter membrane fluidity can disrupt biofilms in the early stages of maturation.

**Figure 4.**
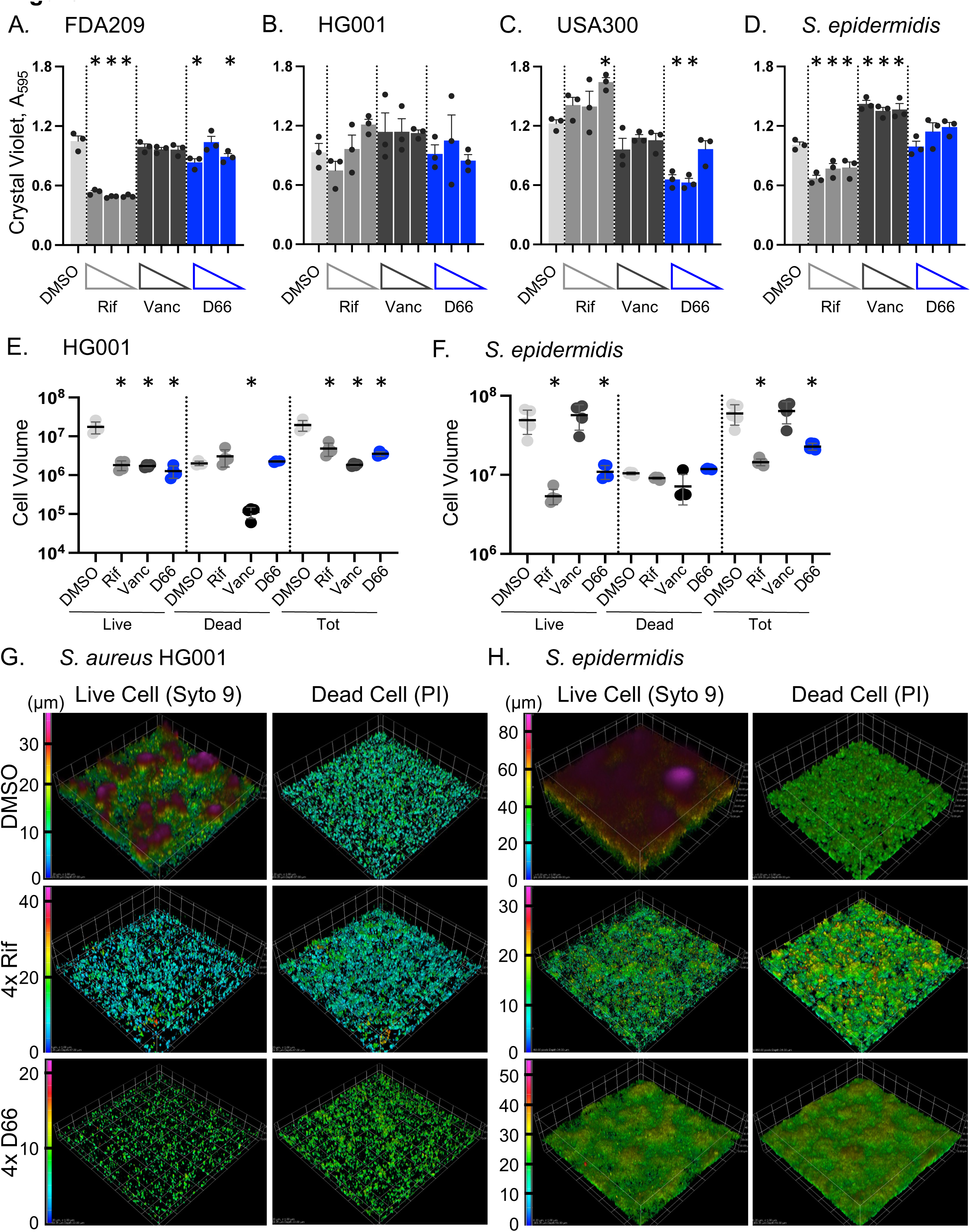
D66 decreases the number of live cells in one-day-old staphylococcal biofilms. For the indicated strain, biofilms established in TSB for 24 hours were treated for 18 hours with DMSO, rifampin (1x MIC_95_ = 0.05 µg/mL), vancomycin (1x MIC_95_ = 1 µg/mL), or D66 (Figure 1B). **A - D)** Remaining biofilm matrix was quantified with crystal violet across 3 compound concentrations (4x, 2x, and 1x MIC from left to right). Mean and SEM from three biological replicates performed in triplicate. Asterisks indicate *P* ≤ 0.05 compared to DMSO, determined by one-way ANOVA with a Dunnett’s post-test. **E, F)** Biofilm volume was measured after staining with Syto9 (live cells) and PI (dead cells). Mean and SD of one of two replicates derived from a minimum of four fields of view. Asterisks indicate *P* ≤ 0.05 compared to DMSO, determined by a one-way ANOVA with a Dunnett’s post-test. **G, H)** Representative Syto 9- and PI-stained biofilm images. Z-stacked images were converted to volumes. Volume data were compiled and reconstructed in Nikon Elements Advanced Research software using a color-coded volume display where blue is the bottom, lowest Z-stack and magenta is the highest point for each sample. Scale bar is in µm. Mean and SD of one of two biological replicates derived from a minimum of four fields of view. Asterisks indicate *P* ≤ 0.01 compared to DMSO, determined by a one-way ANOVA with a Tukey post-test on log_10_ transformed data. G-H are representative images from this same replicate as in E-F.

### Broth persister cells and intracellular S. aureus are sensitive to D66

Staphylococcal biofilms include live cells that are growing and are antibiotic sensitive, in addition to non-growing persister cells that are recalcitrant to antibiotics (4, 36). Since D66 reduced the number of live cells during biofilm formation and early development, we tested whether broth persister cells are susceptible to this compound. Persister cells were generated by growing cultures overnight (37, 38). Samples were monitored by enumeration of CFU after 3 and 24 hours for recalcitrance to high doses of vancomycin (10 μg/mL) and ciprofloxacin (4 μg/mL). In the three *S. aureus* strains, ciprofloxacin prevented growth, an indication of persister cells (39) (**Figure 5A - C**). In the *S. epidermidis* strain, vancomycin prevented growth (**Figure 5 D**). Exposure to D66 reduced survival relative to antibiotic-treated cells in *S. aureus* HG001, indicating that D66 can kill staphylococcal persister cells.

**Figure 5.**
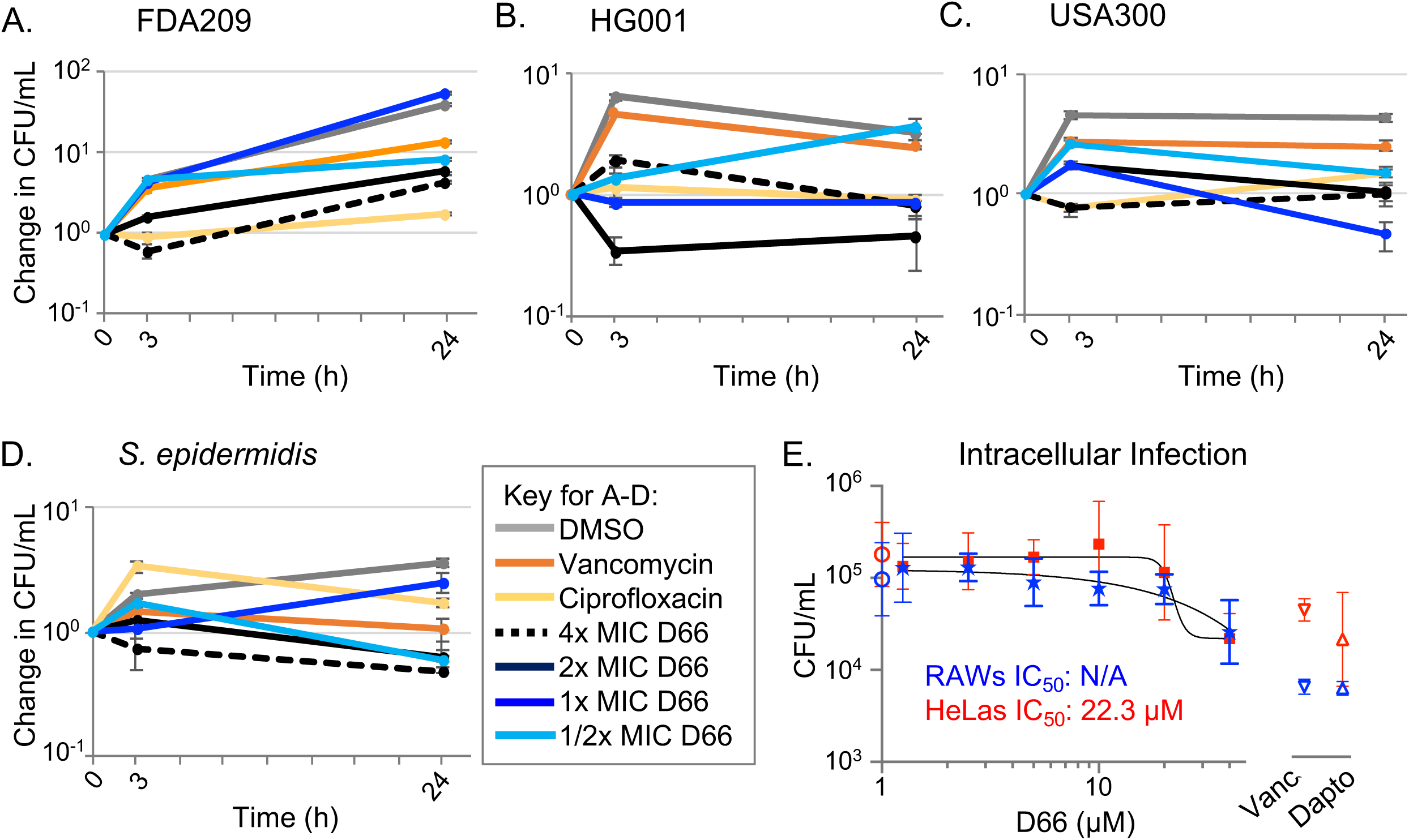
Treatment with D66 reduced the number of viable *S. aureus* broth HG001 persister cells and modestly decreased the load of USA300 in mammalian cells. **A - D)** Overnight cultures of the indicated strains were treated at time 0 with DMSO, ciprofloxacin (4 µg/mL), vancomycin (10 µg/mL), or four different concentrations of D66. Cultures were plated for enumeration of CFU at the time points indicated. Mean and SEM of three biological replicates are shown. **E)** RAW 264.7 macrophage-like cells (blue) and HeLa cells (red) were infected with *S. aureus* USA300, at MOIs of 0.5 and 2.5, respectively. Cells were treated two hours after infection with DMSO (circles on Y axis), vancomycin (vanc, 50 μg/mL, upside down triangles), daptomycin (dapto, 50 μg/mL, right-side up triangles) or dilutions of D66 from 40 µM. After 8 hours of infection, cells were lysed and plated for enumeration of CFU. Mean and SEM of biological duplicates performed with technical triplicates are shown. Nonlinear regression; four parameter variable slope. The IC_50_ values are indicated.

Some *S. aureus* strains can infect and survive within macrophages and epithelial cells, including the USA300 strain (40–43). We therefore tested whether D66 reduces viable intracellular USA300 bacteria in RAW264.7 macrophage-like cells and in HeLa cells. After 8 hours of infection, D66 treatment decreased the number of recoverable bacteria by approximately five-fold; IC_50_ values could be calculated only for *S. aureus* in HeLa cells (**Figure 5E**). The vancomycin and daptomycin controls performed similarly in HeLa cells, and somewhat better in the macrophage-like cells, reducing bacterial load approximately 10-fold. We conclude that D66 and control antibiotics have modest activity against intracellular *S. aureus* survival/growth.

## Discussion

The bacterial cell membrane is an untapped potential source of new antimicrobial targets. While compounds that, like detergents, rapidly permeabilize cell membranes can inhibit persister cells and biofilms (44, 45), whether compounds with more subtle effects on membranes could be prototypes for antibiotics or antibiotic adjuvants has not been well explored. In this study, we used Gram-positive bacteria to examine the mechanism of action and potential utility of a small molecule, D66. This compound was previously demonstrated to be minimally toxic to mammalian cells and mice. D66 inhibits *S.* Typhimurium survival in macrophages and curbs systemic infection in mice. However, D66 has been challenging to study in broth because it has difficulty passing through the Gram-negative bacterial outer membrane barrier and is expelled by efflux pumps (11). We found that in Gram-positive bacteria, the compound prevented regrowth from stationary phase, suggesting that it can reach a bacterial target(s). D66 was bactericidal to planktonic mid-log phase *S. aureus* in broth at concentrations of 1X MIC (80 μM), and the compound fluidized the *S. aureus* cell membrane with a rapidity that suggests fluidization as a primary or near-primary effect of the compound. D66 also modestly disrupted membrane voltage, potentially a secondary effect of membrane fluidization. However, *S. aureus* cells treated with D66 did not rapidly or strongly become permeabilized to large (600 MW) molecules, and the cells maintained a proton gradient, indicating that the compound has a significant but non-catastrophic effect on bacterial cell membrane integrity.

While compounds that lyse membranes disrupt biofilms (46), whether compounds that increase membrane fluidity affect biofilms is not clear. Under growth conditions that promote biofilm formation, D66 had varied and mild effects on the extracellular matrix and on the volume of live and dead cells between strains. In one-day-old biofilms, D66 reduced the volume of live cells as well as rifampin and had a more modest effect on reducing the mass of extracellular matrix. The compound strongly disrupted the structure of one-day-old biofilms, reducing the depth of the mat apparently by reducing the volume and height of the live cell mass. In five-day-old biofilms, D66 exposure reduced biofilm mass only in one staphylococcal strain tested. The negative effects of D66 on biofilms may reflect the membrane fluidizing activity of this compound. Bacterial membrane fluidity and permeability are typically a function of the overall acyl chain composition of the lipids (47). Upon transition from a planktonic to a biofilm growth mode, bacterial cell membranes become increasingly rigid. For example, in *Pseudomonas aeruginosa,* the inner membranes of cells within biofilms are less fluid than those of planktonic bacteria (28, 29). Across Gram-negative and Gram-positive bacteria, cells within biofilms have membranes with a higher percentage of saturated fatty acids than planktonic cells, consistent with increased membrane rigidity, decreased exchange with the environment, and resistance to toxic molecules (30, 31). As bacteria shift from a planktonic to a biofilm growth state, enzymes involved in lipid saturation and membrane rigidification become increasingly expressed (31). Therefore, compounds that enhance membrane fluidity have the potential to interfere with biofilm formation.

D66 exposure reduces staphylococcal persister cell survival in broth and has a modest inhibitory effect on *S. aureus* cells within mammalian cells. These observations suggest that, as in biofilms, bacteria in different growth states germane to infection need to maintain membrane fluidity within bounds to remain viable. Alternatively, or in addition, bacteria in distinct growth states could have varying amounts of a D66 target(s), or enable more D66 to reach a target. Along these lines, persister cells and biofilms are both recalcitrant to antibiotics, as are stationary phase bacteria, in comparison to mid-log phase bacteria, potentially due to differences in membrane permeability (48, 49). In macrophages, D66 could have activity against pathogens within vesicles in part due to lysosomal trapping of the compound with the acidification of the phagosome (13). Phagosomes that harbor *S. aureus* also become acidified, and, as with *S.* Typhimurium, low pH induces virulence gene expression (50–52). However, some *S. aureus* strains escape low pH phagosomes and survive for a period of time within the cytosol (53). We do not yet know whether in macrophages D66 specifically affects *S.* Typhimurium persister cells, growing cells, or both. Given the emerging lifestyle diversity of *S. aureus* strains and *S.* Typhimurium strains within host cells (54, 55), multiple approaches will likely be needed to thwart intracellular infection states across staphylococcal species and strains.

Limitations of this study include the small number of bacterial species in which the effect of D66 on membrane fluidity, permeability, and biofilms were examined. In addition, whether chemical analogs of D66 could separate the effects of the compound on membrane fluidity and growth in broth and in a biofilm state remains to be explored. It will further be of interest to establish whether D66 affects cell membrane fluidity uniformly or in patches throughout the membrane and the cell (56). A more precise understanding of the downstream cellular effects of D66 will be needed to clarify how this compound and possibly others that fluidize membranes act and could be deployed.

Overall, the work highlights that the search for compounds that interfere with biofilms need not be constrained to those that rapidly lyse and/or permeabilize membranes. As existing antibiotics and biocides continue to lose their utility (57), the future of antibacterials and biofilm prevention is likely to require cocktails of functionally complementary agents. It makes sense to include in these cocktails compounds with diverse mechanisms of actions, such as those that modify the fluidity of bacterial cell membranes.

## Methods

### Bacterial Strains

*S. aureus* FDA209 (ATCC 6538), *S. aureus* HG001 (AH2183), *S. aureus* USA300, and *S. epidermidis* 1457 (AH2490). *S. aureus* FDA209 was isolated from a skin lesion, is sensitive to antibiotics, and has historically been used in antimicrobial and quality control testing (***58–60***). *S. aureus* HG001 is a common lab strain derivative of NCTC835, a methicillin-sensitive strain isolated from a septic patient but with a restored *rsbU1* gene to increase its biofilm production (61, 62). *S. aureus* USA300 grows rapidly and is a community-acquired methicillin-resistant strain that causes intracellular infections (62). *S. epidermidis* 1457 is antibiotic sensitive and forms robust biofilms (63, 64). *B. subtilis* 168 is a standard laboratory strain originally isolated from soil (65).

### Media and reagents

Unless otherwise stated, bacteria were grown in lysogeny broth (LB) at 37°C with aeration. D66 (AW00798, MolPort). To obtain mid-log phase cells, bacteria were grown overnight in LB, diluted the next morning 1:100 in fresh LB, and then grown to mid-log phase (OD_600_ 0.4-0.6).

### Minimum inhibitory concentrations

Overnight cultures were diluted in LB to an optical density at 600 nm (OD_600_) of 0.01 and distributed into polystyrene 96-well flat-bottom plates (Greiner, 655185). Compound was added to the desired final concentration, and the final DMSO concentration never exceeded 2%. Plates were grown at 37°C with shaking for 18 hours and OD_600_ was monitored (BioTek Synergy H1 or BioTek Eon). MICs were defined as the concentration at which 95% of growth was inhibited (OD_600_).

### Growth and kill curves

Mid-log phase cultures were sampled at time 0 and then compound or vehicle control (DMSO) was added. Cultures were incubated at 37°C with agitation. At the time intervals indicated, aliquots were monitored for OD_600_ and plated for CFU enumeration. Data for OD_600_ and CFU/mL were normalized to time 0.

### Membrane fluidity

Cells were grown to mid-log phase in LBg (LB with 0.2% glucose). Laurdan (Invitrogen) was added to a final concentration of 10 µM and incubated at 37°C with rotation for 30 minutes. Cells were harvested by centrifugation, washed three times, and resuspended in prewarmed PBSg (PBS with 0.2% glucose). Cells (200 µL) were transferred to a black polystyrene 96-well plate (Greiner, 655076) and monitored (ex 360/em 450 and 500 nm) on a BioTek Synergy H1 plate reader. Baseline fluorescence was recorded for five minutes prior to addition of compound. Fluorescence was recorded for 25 additional minutes. Laurdan generalized polarization (GP) was calculated: GP = (I_460_- I_500_)/ (I_460_+ I_500_).

### Membrane potential

Membrane potential was measured using the potentiometric fluorescent probe DiSC_3_(5) (Invitrogen). Mid-log phase cells were diluted to an OD_600_ of 0.4. DiSC_3_(5) was added to a final concentration of 2 µM and the culture was incubated at 37°C in a rotator for 15 minutes. Cells were captured on a 0.45 µm Metricel^®^ membrane filter (Pall), resuspended in fresh LB, and distributed (200 µL) into black polystyrene 96-well plates (Greiner, 655076). Plates were monitored (ex 650/em 680 nm) on a BioTek Synergy H1 plate reader. After baseline fluorescence was recorded, compound was added to the desired final concentration and measurements were recorded for an additional 30 minutes. The relationship between membrane potential and DiSC_3_(5) fluorescence intensity (calibration) was established as described (23, 66). Briefly, *S. aureus* FDA209 was grown overnight in LB media with 50 mM Hepes pH 7.5 and three concentrations of KCl (46 mM, 18.1 mM and 7.1 mM), with three corresponding concentrations NaCl to maintain ionic strength (254 mM, 281.9 mM, 292.9 mM, respectively). Cells were diluted 1:100 in these same media and grown until mid-log phase. Cells were then diluted to an OD_600_ of 0.3 and incubated with 2 μM DiSC3(5) for 15 minutes. Valinomycin (K+ carrier) was added to a final concentration of 5 μM and fluorescence intensity was monitored with a BioTek Synergy H1 (ex 650/em 680 nm) for 30 minutes.

### Cytosolic pH

BCECF (2’,7’-Bis-(2-Carboxyethyl)-5-(and-6)-Carboxyfluorescein, Acetoxymethyl Ester (BCECF-AM) (Molecular Probes) was added to mid-log phase cells in BCECF LB (LB, 0.1% glucose, 300 mM KCl, 50 mM HEPES, pH 7) to a final concentration of 10 µM and incubated at 37°C in a rotator for one hour. Cells were diluted 1:10 and pipetted into a black polystyrene 96-well plate (Greiner, 655076). After five minutes of equilibration, DMSO, nigericin (Sigma-Aldrich, N7143) or D66 were added and fluorescence (ex 490/em 535 nm and ex 440/em 535 nm) was monitored every 2.5 minutes for 20 minutes using a BioTek Synergy H1 plate reader. BCECF fluorescence was calibrated at 7 pHs between 5.5 and 8 (every 0.5 pH): pH = pK_a_ – log(I_490_/I_440_); the pK_a_ of BCECF is 6.97 (12).

### Reduction potential

Mid-log-phase cells (200 ml per well) were transferred to a black polystyrene 96-well plate (Greiner; 655076) containing compound. Compound was added, and the plate was incubated with shaking. Resazurin (alamarBlue; Invitrogen) (67) was added to a final concentration of 100 mg/ml 5 min at the indicated time point. The plate was incubated with shaking in the dark at room temperature for 5 min. Fluorescence readings were taken (ex 570/em 650 nm) using a BioTek Synergy H1 plate reader.

### Cytosolic ATP

Intracellular ATP levels were measured using BacTiter-Glo Microbial Cell Viability Assay (Promega) according to the manufacturer’s instructions. Overnight cultures were sub cultured in fresh LB and grown to an OD_600_ of 0.35 - 0.45. Cells (100 μL) were added to 2 μL of compound in a 96-well plate and incubated for 10 minutes at 37 °C with agitation. Reagent (100 μL) was added and incubated in the dark with agitation for 5 minutes. Luminescence was read on a BioTek Synergy H1 plate reader.

### Membrane barrier assays

Compound, DMSO, or SDS as stated on figure was added to mid-log phase cells to the desired concentration, and cultures were sampled at 0, 10, 15, 20, 30 and 45 minutes. Five minutes before harvesting, PI [10 µg/mL] (Life Technologies) was added. Cells were pelleted, washed twice, resuspended in PBS, and monitored (ex 535/em 617 nm) using a BioTek Synergy H1 plate reader.

### Single-cell membrane potential and permeability

Overnight cultures of *S. aureus* strain FDA 209 were diluted 1:100 in fresh LB and incubated until grown to mid-log phase (OD_600_ 0.3 - 0.6). Cultures were treated with varying concentrations of D66, JD1, Nisin, CCCP or DMSO and incubated at 37°C rotating for either 5 minutes, 20 minutes, or 45 minutes. 30 μM DiOC2(3) and 200 nM TO-PRO-3 were then added, and the cultures were incubated in the dark at 37°C rotating for an additional 15 minutes. Samples were run using the Beckman Coulter CyAn ADP Analyzer with Summit version 4.4. Bacteria were identified based on SSC versus FSC. DiOC2(3) (ex 488 nm/ em PMT FL1; green, and FL3; red) and TO-PRO-3 (ex 635 nm/em PMT FL8) were quantified across at least 40,000 events per sample. Data was analyzed using FlowJo version 10 9.0.

### Biofilm growth and treatment

As previously described (27), cultures were grown to mid-log phase in TSB, diluted to 4 x 10^6^ CFU/mL in TSB and 200 µL were added to each well of a flat-bottomed, polystyrene, 96-well (Greiner 655185) plate (edge wells on the plate were filled with PBS to minimize evaporation of experimental wells). After 24 hours of incubation at 37°C without agitation wells were carefully washed twice with PBS followed by the addition of 200 µL of TSB containing 4 µL DMSO, or antibiotic or compound to achieve the desired final concentration. For 5-day biofilm assays wells were washed and received fresh media daily and DMSO, antibiotic, or compound were added on day 5. Plates were incubated at 37°C for 18 hours without agitation. Prior to staining, the biofilm was washed twice with PBS to remove non-adherent cells.

### Biofilm inhibition assays

Cultures were grown to mid-log phase in TSB, diluted to 4 x 10^6^ CFU/mL in TSB, and 200 µL of this dilution was added to each well of a flat-bottomed, polystyrene, 96-well (Greiner 655185) plate, and 4 µL of DMSO, antibiotic, or compound was added to achieve the desired final concentration. Edge wells on the plate were filled with PBS to minimize evaporation of experimental wells. After 24 hours of incubation at 37°C with no agitation the media and planktonic cells were carefully removed, and wells were washed 2 times with PBS. A multichannel pipette was used for removal of media and subsequent PBS washes.

### Biofilm mass assessment with crystal violet

Plates containing washed biofilms were allowed to air dry then 200 µL of 0.01% crystal violet was added to each well and incubated for 20 minutes. The biofilms were washed once with DI water and air dried. The stained biofilm was resuspended in 200 µL 70% ethanol and the A_595_ was measured.

### Biofilm volume microscopy and analysis

Biofilms were grown as described above with the following differences (27). Cellvis 35mm # 1.5H Glass Bottom Dish with 20 mm bottom well dishes were used for biofilm growth and PBS washes to remove non-adherent cells (cells not attached to the biofilm) were performed with serological pipettes to minimize disruption. Biofilms were stained for 15 minutes prior to imaging with PI [10 µg/mL] and Syto9 [3 µM] resuspended in PBS. The dye solution was gently removed with a pipette, and samples were washed twice with PBS and imaged live (fixation with 4% PFA resulted in a PI-staining artifact). Images were acquired using the Yokogawa CellVoyager CV1000 Confocal Scanner System with a 40x/0.6NA WD 2.7-4.0 (mm) objective, a Microlens-enhanced Nipkow Disk Scanner with a pinhole size of 50 um and Hamamatsu Photonics | ImagEM X2 EM-CCD Camera C9100-14 High Resolution 16-bit format. Prior to image acquisition, each biofilm Z range was determined by manually establishing the top and bottom of the biofilm, and the z-step size was set to 1 µm to simplify volume analysis (conversion between volume and voxels). Images were acquired in two color channels: ex/em for Syto 9 and PI at 488nm_525/50 and 561nm_617/73, respectively. A minimum of eight randomly selected fields of view were acquired and at least four images were used for analysis. Images were imported to MATLAB R2020b as multidimensional tiff stacks and processed as volume data using a custom script designed to store all results in data structures for user review. To create 3D binary masks for extracting the Syto 9 and PI foreground signals from the background Otsu’s threshold was chosen over manual thresholding due to widely differing levels of background signal observed across strains and/or caused by biofilm thicknes*s* (68). The total volume for each of the two channels were quantified as the summation of voxels identified above the threshold per channel. The total number of voxels were converted to total volume (1 voxel = 0.0367 µm^3^) using post analysis of metadata. Select volumes were reconstructed in Nikon Elements using thresholds determined by Otsu’s method to provide a color-coded quantitative volume display based on depth in micrometers. The DMSO, rifampin and vancomycin control samples were from the same experiments as the D66-treated samples but were published previously (27).

### Persister assays

Cultures were grown overnight in Trypticase soy broth (TSB) at 37°C with aeration for 18 h. Cultures were divided into samples and antibiotic or compound was added to the indicated final concentration. Samples were incubated at 37°C with aeration and plated at 0, 3, and 24 h for enumeration of CFU.

### Intracellular infection assays

RAW 264.7 (TIB-71) macrophages (5 x 10^4^ macrophages in 100 μL of complete DMEM) were seeded in 96-well tissue culture plates (Greiner, 655180) and were incubated at 37°C with 5% CO_2_. For experiments performed with HeLa cells (ATCC CCL-2), 1 x 10^4^ cells were seeded per well. *S. aureus* USA300 was grown overnight in TSB and sub-cultured in TSB to an OD_600_ of 0.4 for regrowth to an OD_600_ 0.6. Cultures were diluted into to a final concentration of 5 x 10^5^ CFU/mL in complete DMEM. Twenty-four hours after seeding of the mammalian cells, 50 µL of bacterial cultures were added to each cell culture well, at approximate multiplicities of infection (MOI) of 0.5 and 2.5, respectively, for RAW 264.7 cells and HeLa cells. Plates were centrifuged at 500 *x g* for 2 minutes to synchronize infection. Thirty or forty-five minutes (for RAW 264.7 or HeLa cells, respectively) after bacterial addition, wells were washed one time with PBS. DMEM containing 100 µg/mL gentamicin (Sigma-Aldrich) was added and cells were incubated for 90 minutes. Wells were then washed twice with PBS. Complete DMEM was added to the wells followed by 1 μL of DMSO or D66 to the stated final concentration. After 8 hours of infection, cells were lysed and plated for CFU.

### Statistical Analyses

GraphPad Prism version 10.11 was used for statistical comparisons.

## Acknowledgements

We thank all the members of the Detweiler laboratory for insightful discussions and technical help. We are grateful to C.K. Asamoto, C.A. Ewing, D. Jiang, and C.T. Meyers for insightful comments on the manuscript. We thank the MCDB Light Microscopy Facility at the University of Colorado Boulder.

---

**Figure S1.**
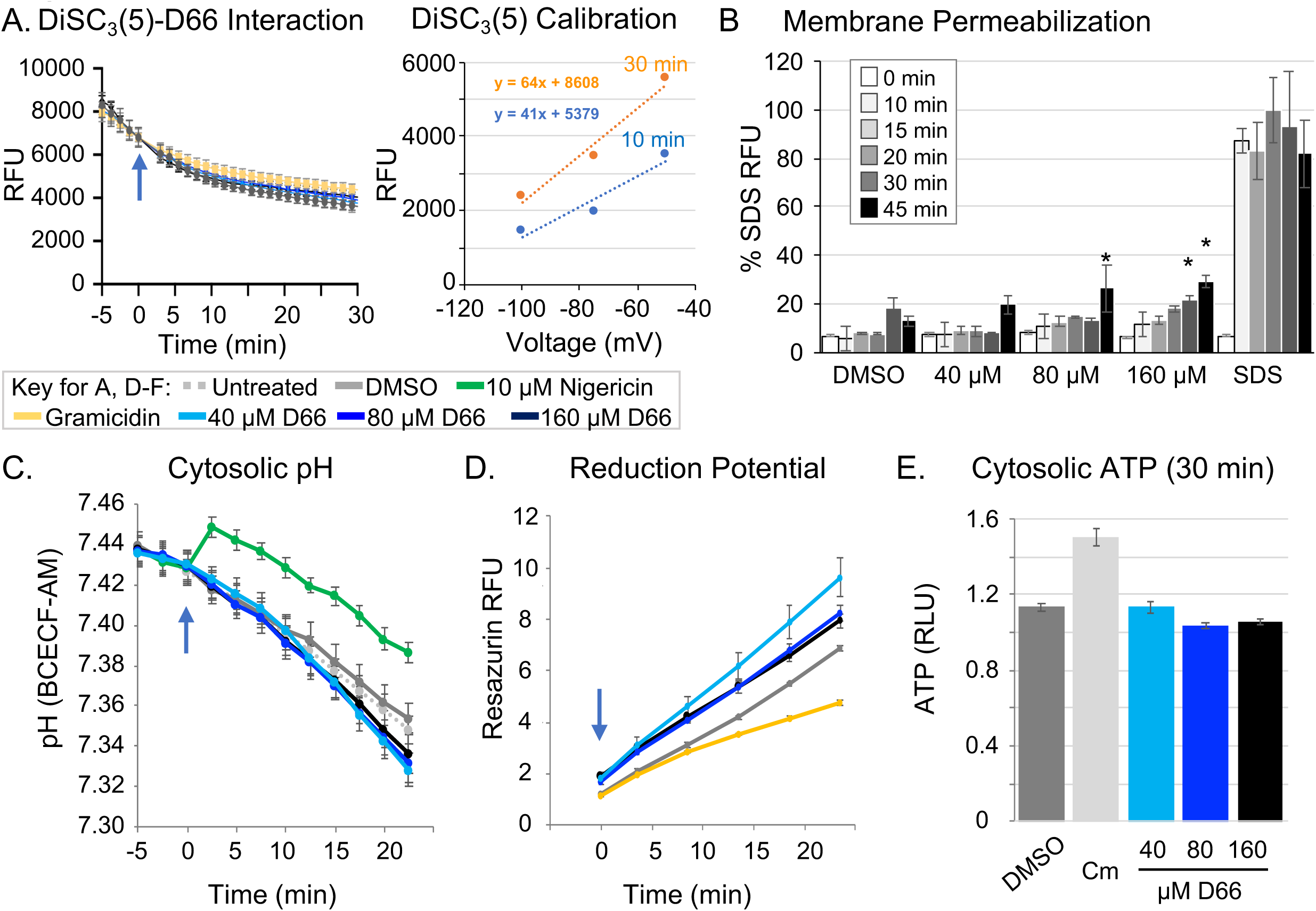
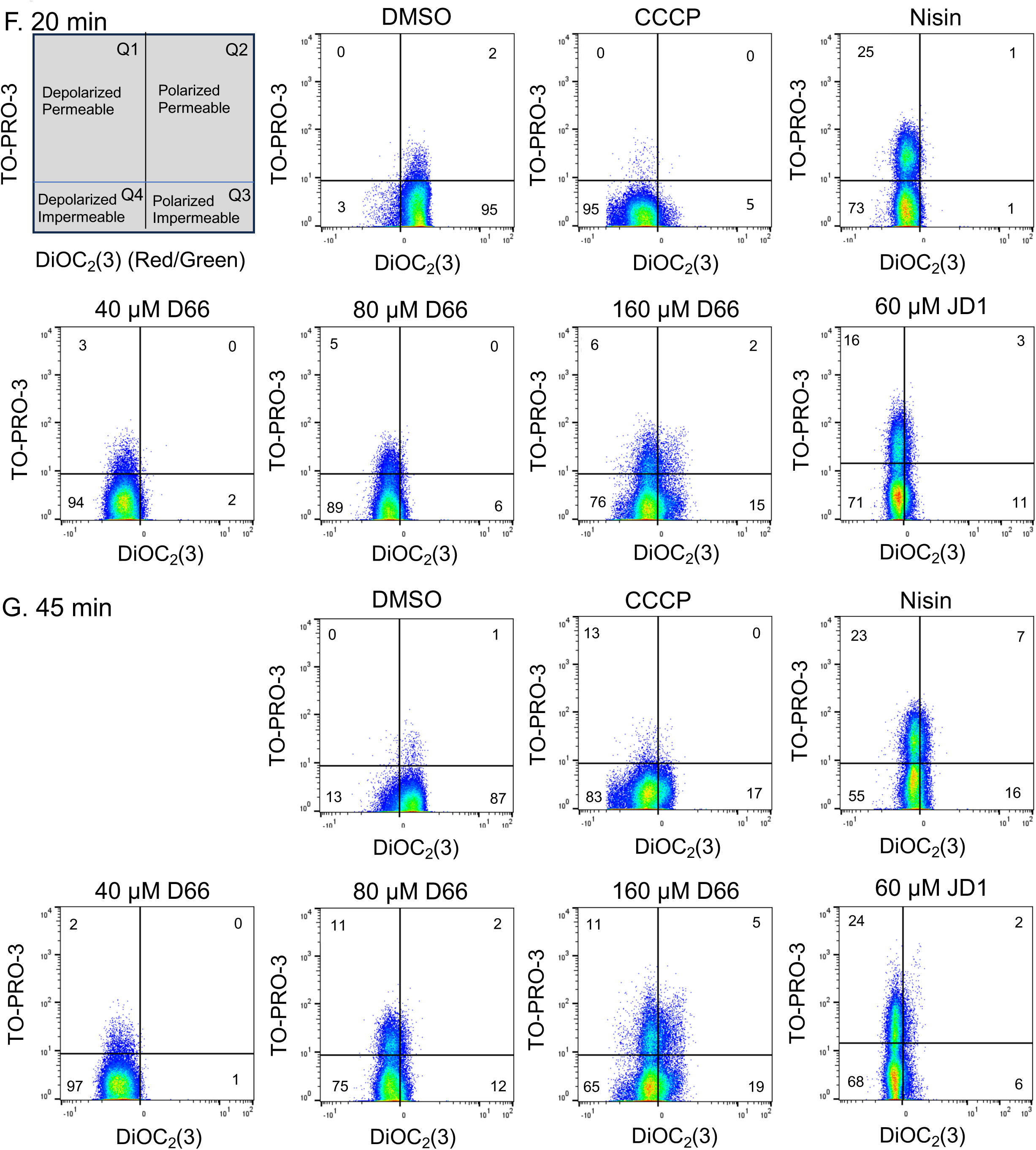
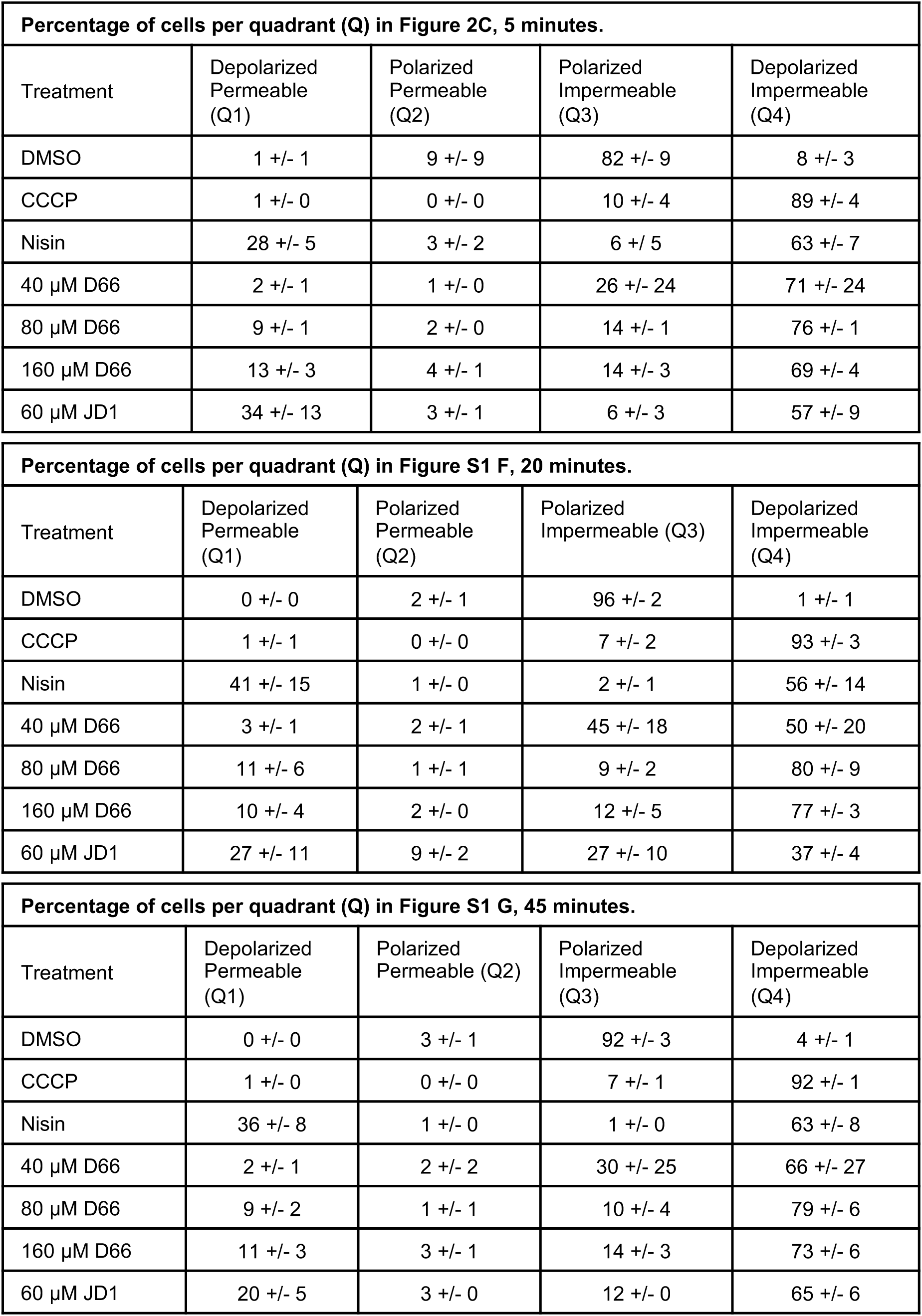
Supporting data for Figure 2. Mid-log phase *S. aureus* FDA209 cells were used for all experiments. **A)** DiSC_3_(5) interaction with D66 (left) and calibration of DiSC_3_(5) RFU for FDA209. D66 or gramicidin (2 μg/ml; 1.1 mM) were added to DiSC_3_(5) just after time 0 (arrow). Data were normalized to DMSO at time 0. Relative fluorescent units (RFU). Mean and SEM of three biological replicates performed with technical triplicates. Calibration of DiSC_3_(5) RFU was based on potassium concentrations in the medium (see methods). The 10- and 30-minute post-treatment timepoints correspond to those in Figures S1A and 2B, C. **B)** Membrane permeability was monitored by PI fluorescence. Cells were treated just after time 0 with DMSO, SDS (0.3%), or D66. Samples were processed at the timepoints shown. Mean and SEM of three biological replicates performed with technical triplicates, normalized to the highest SDS value (20-min). Asterisks indicate *P* ≤ 0.05 compared to DMSO, determined by a one-way ANOVA with a Tukey post-test. **C)** Intracellular pH was measured with the fluorescent probe BCECF-AM. Cells were treated just after time 0 (arrow) with DMSO, the protonophore nigericin [10 µM], or D66. Mean and SEM of three biological replicates performed with technical triplicates. **D)** Reduction potential (respiration) was determined upon incubation with resazurin (alamar blue) and treatment just after time 0 (arrow). Mean and SEM of three biological replicates performed with technical triplicates and normalized to untreated cells. **E)** Membrane permeability, as monitored by ATP leakage. with the Promega BacTiter-Glo kit after 30 minutes of treatment with DMSO, chloramphenicol (Cm) (32 mg/mL), or D66. Mean and SEM of relative luciferase units (RLU) for three biological replicates performed with technical triplicates and normalized to untreated cells. **F, G)** Membrane polarization and permeability were determined by flow cytometry using the fluorescent dyes TO-PRO-3 and DiOC_2_(3). Cells were treated with DMSO, CCCP [30 μM], nisin [75 μg/mL] or D66 for 20 or 45 minutes. Data shown are representative of three biological replicates. **H)** Mean and SEM of the flow cytometry data at 5, 20, and 45 minutes.

**Figure S2.**
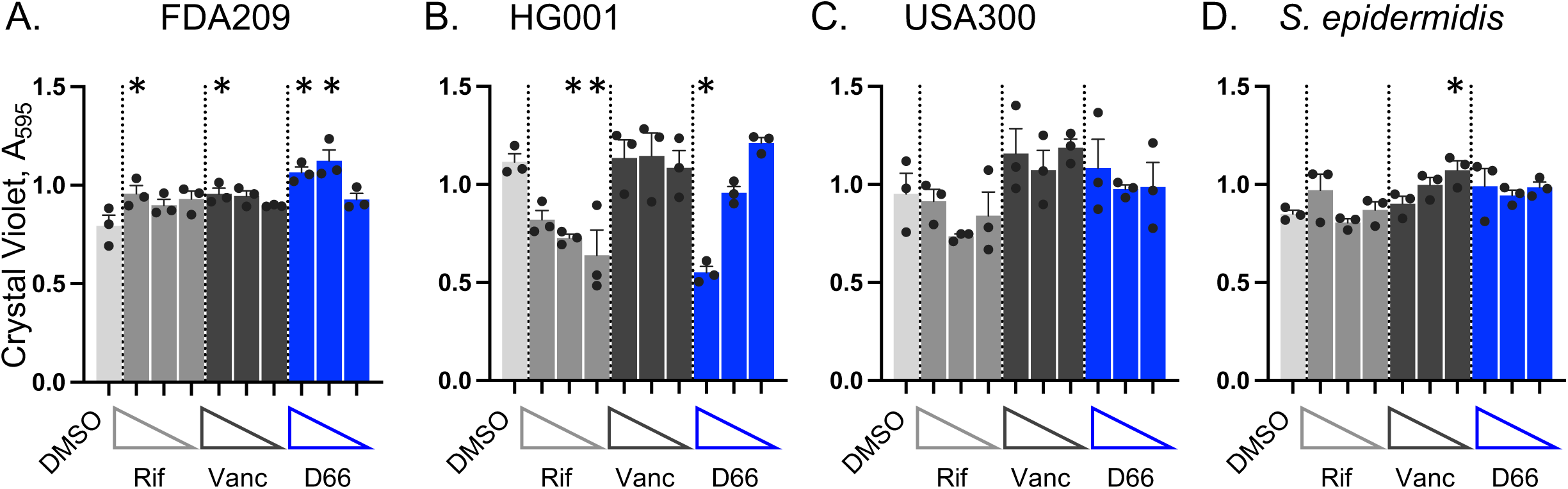
D66 minimally affects 5-day-old staphylococcal biofilms. A-D). For the indicated strain, biofilms established in TSB for 5-days were treated for 18 hours with DMSO, rifampin (1x MIC_95_ = 0.05 µg/mL), vancomycin (1x MIC_95_ = 1 µg/mL), or D66 (Figure 1B). Remaining biofilm matrix was quantified with crystal violet across 3 compound concentrations (4x, 2x, and 1x MIC from left to right). Mean and SEM from three biological replicates performed in triplicate. Asterisks indicate *P* ≤ 0.05 compared to DMSO, determined by one-way ANOVA with a Dunnett’s post-test.

